# Cytoplasmic domain and enzymatic activity of ACE2 is not required for PI4KB dependent endocytosis entry of SARS-CoV-2 into host cells

**DOI:** 10.1101/2021.03.01.433503

**Authors:** Hang Yang, Xiaohui Zhao, Meng Xun, Lingjie Xu, Bing Liu, Hongliang Wang

## Abstract

The recent COVID-19 pandemic poses a global health emergency. Cellular entry of the causative agent SARS-CoV-2 is mediated by its spike protein interacting with cellular receptor- human angiotensin converting enzyme 2 (ACE2). Here, we used lentivirus based pseudotypes bearing spike protein to demonstrate that entry of SARS-CoV-2 into host cells is dependent on clathrin-mediated endocytosis, and phosphoinositides play essential role during this process. In addition, we showed that the intracellular domain and the catalytic activity of ACE2 is not required for efficient virus entry. These results provide new insights into SARS-CoV-2 cellular entry and present potential targets for drug development.

## Introduction

Since the beginning of this century, there have been three beta-coronaviruses crossed the species barrier to cause severe respiratory diseases. In 2002-2003, sever acute respiratory syndrome coronavirus (SARS-CoV) infected 8096 people and caused 774 deaths(1). In 2012, Middle East respiratory syndrome coronavirus (MERS-CoV) was discovered as causative agent of a severe respiratory syndrome in the middle east area and as of January 2020, more than 2500 people was diagnosed with MERS, with 866 associated deaths (WHO). In late 2019, a novel coronavirus, named SARS-CoV-2 emerged and soon transmitted to cause a globally pandemic. As of January 2021, there were over 100 million confirmed cases of COVID-19 worldwide, resulting in more than 2 million deaths (coronavirus.jhu.edu). All three coronaviruses are believed to originate from bats but zoonotic transmission involved different intermediate hosts. With numerous coronaviruses in bats, it is likely that coronavirus crises will continue to occur in the foreseeable future (2).

Cellular entry of coronavirus is mediated by its spike glycoprotein (S) by interacting with cellular receptors. The sequences of SARS-CoV and SARS-CoV-2 S protein are conserved and numerous studies have shown that they exploited the same receptor, angiotensin-converting enzyme 2 (ACE2)(3–6). Different from SARS-CoV, the S protein of SARS-CoV-2 was efficiently processed into two subunits, S1 and S2, which mediates attachment and membrane fusion, respectively(3, 6). Structural analysis showed that S protein formed trimer with the receptor-binding domain exposed up for easy receptor accessibility(5, 6). ACE2 is a type I transmembrane protein with an extracellular domain with homology to ACE and a short cytoplasmic tail(7, 8). ACE2 is a Zinc metalloprotease that catalyzes the cleavage of Ang I to Ang1-9, but its catalytic activity is not required for spike induced syncytia formation(9).

The entry of enveloped viruses into cells is known to occur via two primary pathways, i.e. direct membrane fusion at the cell surface, or endocytosis, with the latter being a pH-sensitive process(10). The endocytic pathways exploited by animal viruses to gain entry into host cells include clathrin-mediated endocytosis (CME), caveolae-dependent endocytosis, as well as poorly characterized routes such as clathrin- and caveolae-independent endocytosis(11). SARS-CoV and MERS-CoV S can be triggered to fuse either at the plasma membrane or the endosomal membrane depending on protease availability(12). In addition, SARS-CoV has been shown to enter host cells via direct membrane fusion, clathrin-mediated endocytosis or non-clathrin, non-caveolae-mediated endocytosis(13–15), suggesting the virus might employ various strategies to expand its cellular tropism. The specific endocytic pathways SARS-CoV-2 exploited have not been fully characterized.

Phosphoinositides(PI) are known to be involved in the whole process of endocytosis(16), of note, PI(4,5)P_2_ has long been thought to be the most important PI during CME (17–21). A recent study demonstrated that a successive PI molecular conversions accompanies the clathrin-coated pit assembly, budding and uncoating with PI(4,5)P_2_, PI4P, PI(3,4)P_2_ function at different stages of CME(22). Interestingly, SARS-CoV has been reported to require PI4KIIIβ for cell entry and depletion of PI4P by Sac1 inhibited viral entry(23), suggesting a role of PI4P during viral entry. In contrast, inhibitors targeting PI(3,5)P_2_ blocked SARS-CoV-2 viral entry (4). Whether other PI molecules are required for SARS-CoV-2 entry is still unknown.

Here we demonstrated that SARS-CoV-2 entered cells via clathrin-mediated endocytosis, but independent of caveolae-mediated endocytosis. Distinct from SARS-CoV, SARS-CoV-2 did not seem to require lipid rafts for infection. In addition, phosphatidylinositol molecules PI4P and PI(4,5)P_2_ also played essential roles during virus entry. Finally, we showed that the cytoplasmic domain and the enzymatic activity of ACE2 were not required for SARS-CoV-2 entry. Our data provided new insights into SARS-CoV-2 cellular entry and presented potential targets for drug development.

## Results

### Incorporation of S protein into pseudovirus and infection of pseudovirus on different cells

Pseudovirus has been widely used to mimic the entry of real virus and is a powerful tool for studying early events in the life cycle of a virus. We here employed a lentivirus-based pseudovirus to study the entry of SARS-CoV-2. To facilitate the incorporation of SARS-CoV-2 S protein into pseudovirus, we employed a method similar as previously reported in(4). A codon-optimized cDNA encoding the S protein was synthesized and the C-terminal last 19 amino acids, which contains an endoplasmic reticulum retention signal was replaced with FLAG tag (Fig S1A). Immunoblotting showed that codon-optimized S cDNA expressed at a much higher level than that of native spike in 293T cells (Fig S1B). In addition, consistent with previous reports (3, 4, 6), the S protein was processed and two major bands were observed, reflecting the full-length and cleaved S proteins respectively(Fig S1B). To confirm S protein was efficiently incorporated into the pseudovirus, spike and VSV-G protein-bearing pseudoviruses were pelleted by ultracentrifugation and tested against anti-spike antibody and anti-HIV-p24 antibody (Fig 1A). The results showed that the majority of S proteins on pseudovirus were cleaved, suggesting the S protein was efficiently processed in host cells.

**Fig 1.**
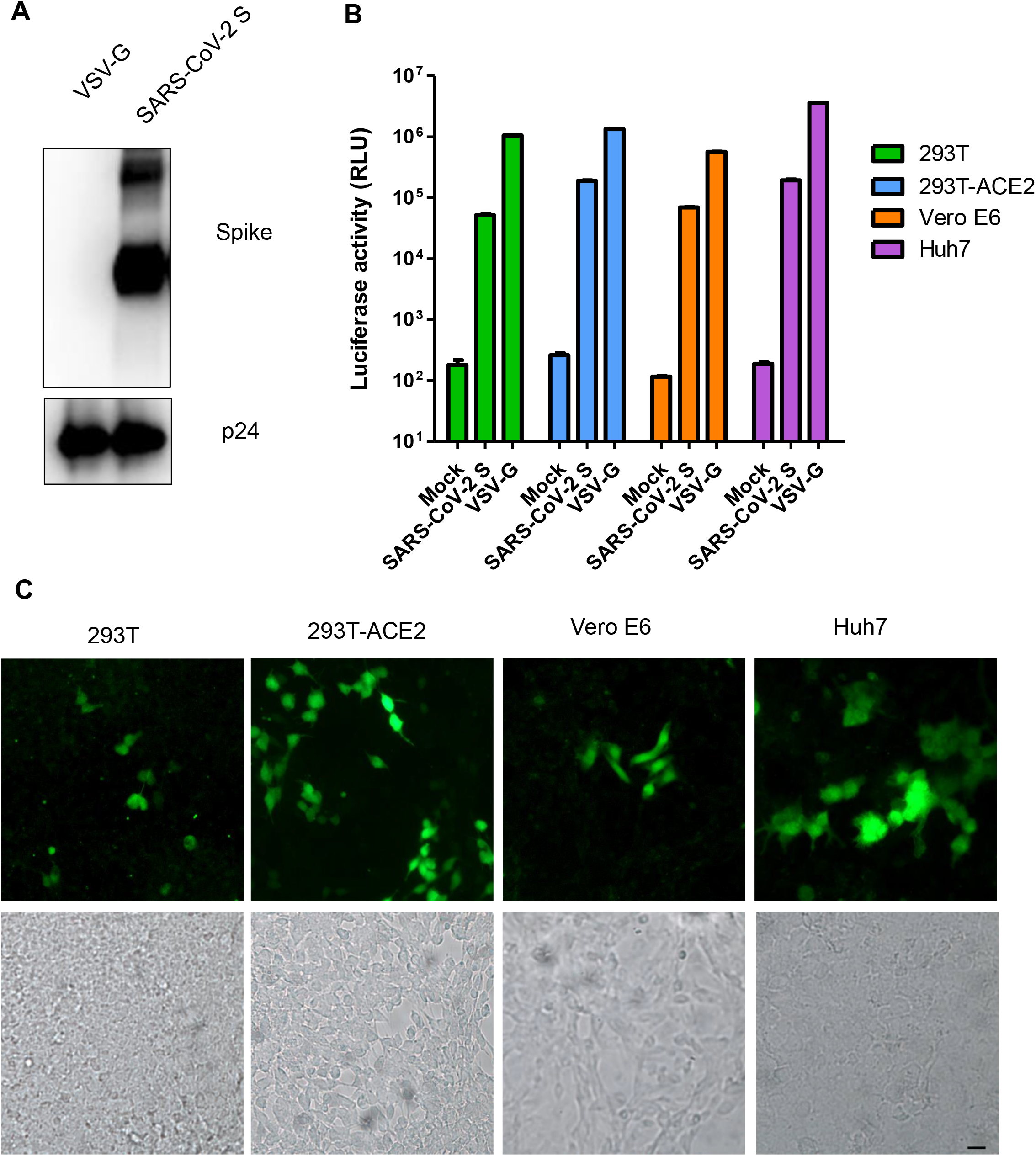
Characterization of SARS-CoV-2 Spike pseudovirus. A. Incorporation of SARS-CoV-2 S protein into pseudoviruses. VSV-G and S pseudoviruses were pelleted as described in Materials and Methods and subjected to SDS-PAGE, immunoblotted with anti-Spike and HIV p24 as control. B. S pseudovirus transduction of 293T, 293T-ACE2, Vero E6 and Huh 7 cells. 293T, 293T-ACE2, Vero E6 and Huh 7 cells were mock infected or infected with S pseudovirus, VSV-G pseudovirus expressing NanoLuc luciferase. At 48 h post transduction, luciferase activity was measured. Values are mean ± SD and are representative of three independent experiments. C. 293T, 293T-ACE2, Vero E6 and Huh 7 cells transduced with S pseudovirus expressing GFP. Bright field was included to show the presence of cells. Bar, 20μm.

Next, we tested whether these Spike-bearing pseudoviruses (S pseudovirus) were able to transduce host cells. For this purpose, we infected 293T, Vero E6, Huh 7 and 293T cells overexpressing ACE2 with S or VSV-G protein pseudovirus containing a luciferase reporter gene. As expected, VSV-G pseudovirus transduced all four cell types efficiently. Compared to 293T cells, overexpression of ACE2 enhanced S pseudovirus infection, suggesting SARS-CoV-2 utilized ACE2 as a receptor. In addition, although both Vero E6 and Huh 7 cells can be transduced by S pseudovirus, Huh 7 gave more than two-fold increase in luciferase activity than Vero E6 cells (Figure 1B). Thus, Huh 7 was chosen for the following viral entry studies. Similarly, GFP-expressing pseudoviruses also transduced the above cells lines successfully (Fig 1C), further suggesting that the S pseudovirus we prepared can be used for viral entry studies.

### S pseudovirus enters cells via clathrin-mediated endocytosis

We next tested whether SARS-CoV-2 enters cells via endocytosis or direct membrane fusion at plasma membrane. In contrast to direct membrane fusion, endocytosis is thought to be pH-dependent(24). When Huh 7 cells were pretreated with lysosomotropic agents, like chloroquine, ammonium chloride or bafilomycin A1, significant decreases of transduction were observed for both VSV-G and SARS-CoV-2 S pseudovirus (Fig 2A), suggesting that SARS-CoV-2 entry of Huh 7 cells is pH-dependent. Consistent with this, dual immunostaining of S protein and early endosome marker EEA1 showed colocalization (Fig 2B), supporting that S pseudovirus infected Huh 7 cells via endocytosis. Importantly, we observed that S pseudovirus transduction correlated with higher EEA1 expression in cells (Fig. S2) and immunoblotting showed that S pseudovirus transduction up-regulated EEA1 expression in Huh 7 cells (Fig 2B). All these suggested that SARS-CoV-2 enters cells via pH-dependent endocytosis.

**Fig 2.**
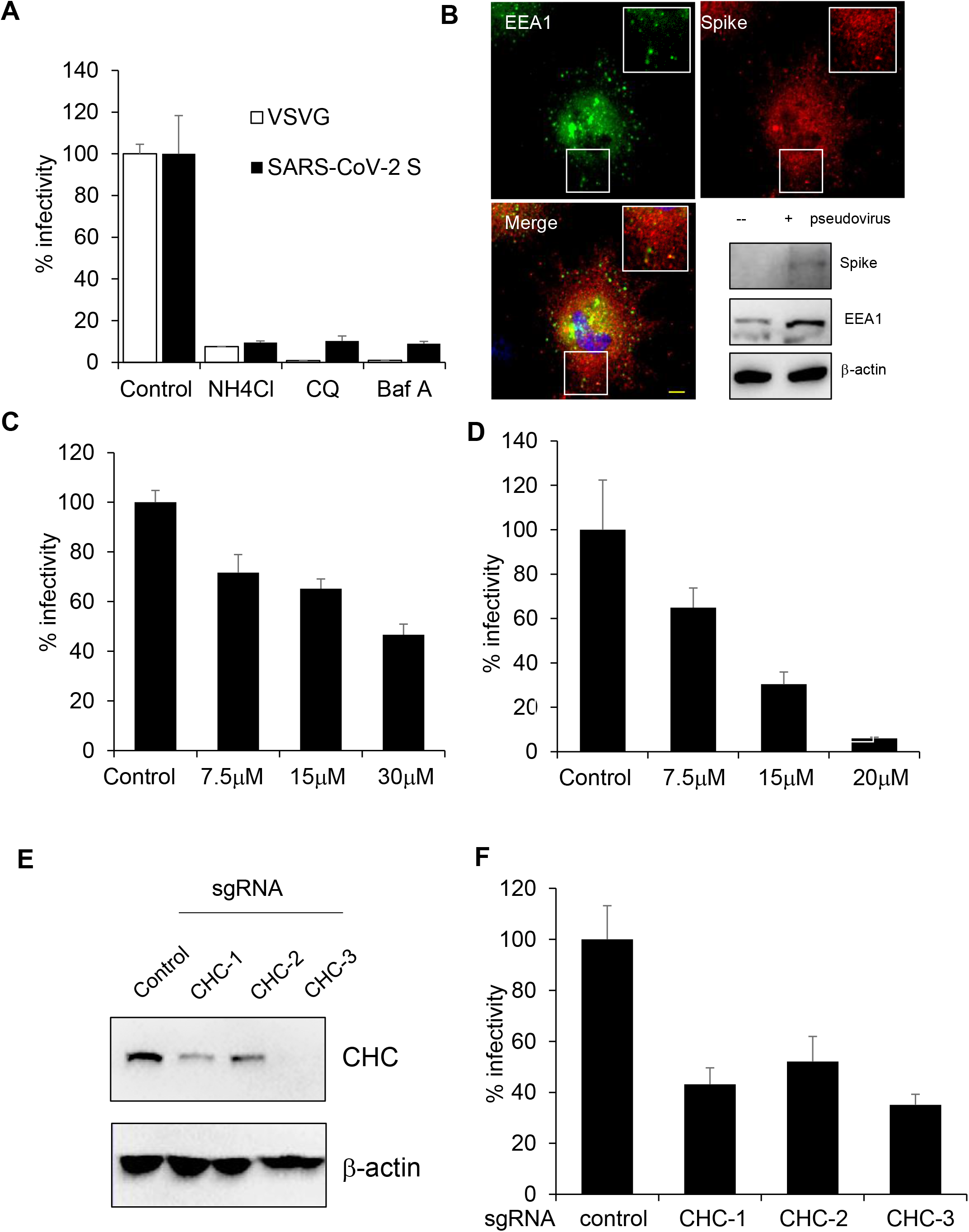
S pseudovirus transduces Huh 7 cells via clathrin-mediated endocytosis. A. Huh 7 cells pretreated with lysosomotropic agents (50mM NH4Cl, 100μM chloroquine and 100 nM bafilomycin A) were transduced with S pseudovirus or VSV-G pseudovirus. Luciferase were measured to reflect viral infection. Values are normalized to control and expressed as mean ± SD. B. Immunostaining of S pseudovirus infected Huh 7 cells for Spike(*red*) and EEA1(*green*). Bar, 10μm. Lower right corner, immunoblotting of EEA1 expression with or without S pseudovirus infection. C. Huh 7 cells pretreated with indicated concentrations of Pitstop were transduced with S pseudovirus and luciferase were measured. Values are normalized to control and expressed as mean ± SD. D. Huh 7 cells pretreated with indicated concentrations of CPZ were transduced with S pseudovirus and luciferase were measured. Values are normalized to control and expressed as mean ± SD. E. Immunoblotting of CHC expression in Huh 7 cells stably expressing a negative control or three independent CHC sgRNAs with Cas9. F. Huh 7 cells as treated in (E) were transduced with S pseudovirus and luciferase were measured. Values are normalized to control and expressed as mean ± SD.

Clathrin-mediated endocytosis is the best documented mode of endocytosis, to test whether SARS-CoV-2 gain entry into cells via CME, we first pretreated cells with clathrin inhibitor, pitstop(25). Pitstop inhibited the entry of SARS-CoV-2 in Huh 7 cells in a dose-dependent manner (Fig 2C). Similarly, chlorpromazine(CPZ), which prevents the assembly of coated pits at the cell surface, also inhibited the entry of SARS-CoV-2 in a dose-dependent manner (Fig 2D). To overcome any nonspecific effect the chemical compounds may have, we also carried out an orthogonal CRISPR/Cas9 knockout experiment. Huh 7 cells stably transduced with lentiviral vectors encoding the sgRNA targeting clathrin heavy chain (CHC) and *Streptococcus pyogenes* Cas9 were first validated of gene targeting (Fig 2E) and then transduced with luciferase encoding S pseudovirus. The magnitude of luciferase activity correlated with the expression of CHC (Fig 2F), suggesting SARS-CoV-2 entry is dependent on clathrin.

### S pseudovirus infection does not require caveolae-mediated endocytosis

Caveolae are small, flask-shaped invaginations in the plasma membrane composed of high levels of cholesterol and glycosphingolipids as well as the integral membrane protein caveolin(26). To determine whether SARS-CoV-2 enters cells through a caveolae-mediated pathway, we first treated cells with filipin or methyl-β-cyclodextrin (MβCD), both of which could disrupt the integrity of cholesterol-enriched membrane microdomains(27). Compared to control cells, uptake of cholera toxin subunit B (CTB), which has been reported to be internalized via caveolae-mediated endocytosis (28–30), was blocked at cell surface when cells were treated with filipin or MβCD (Fig S2), suggesting these drugs can block caveolae-mediated endocytosis. However, neither drug inhibited the huh 7 transduction of S pseudovirus (Fig 3A, 3B), indicating that SARS-CoV-2 entry of Huh 7 cells was independent of cholesterol-enriched membrane microdomains.

**Fig 3.**
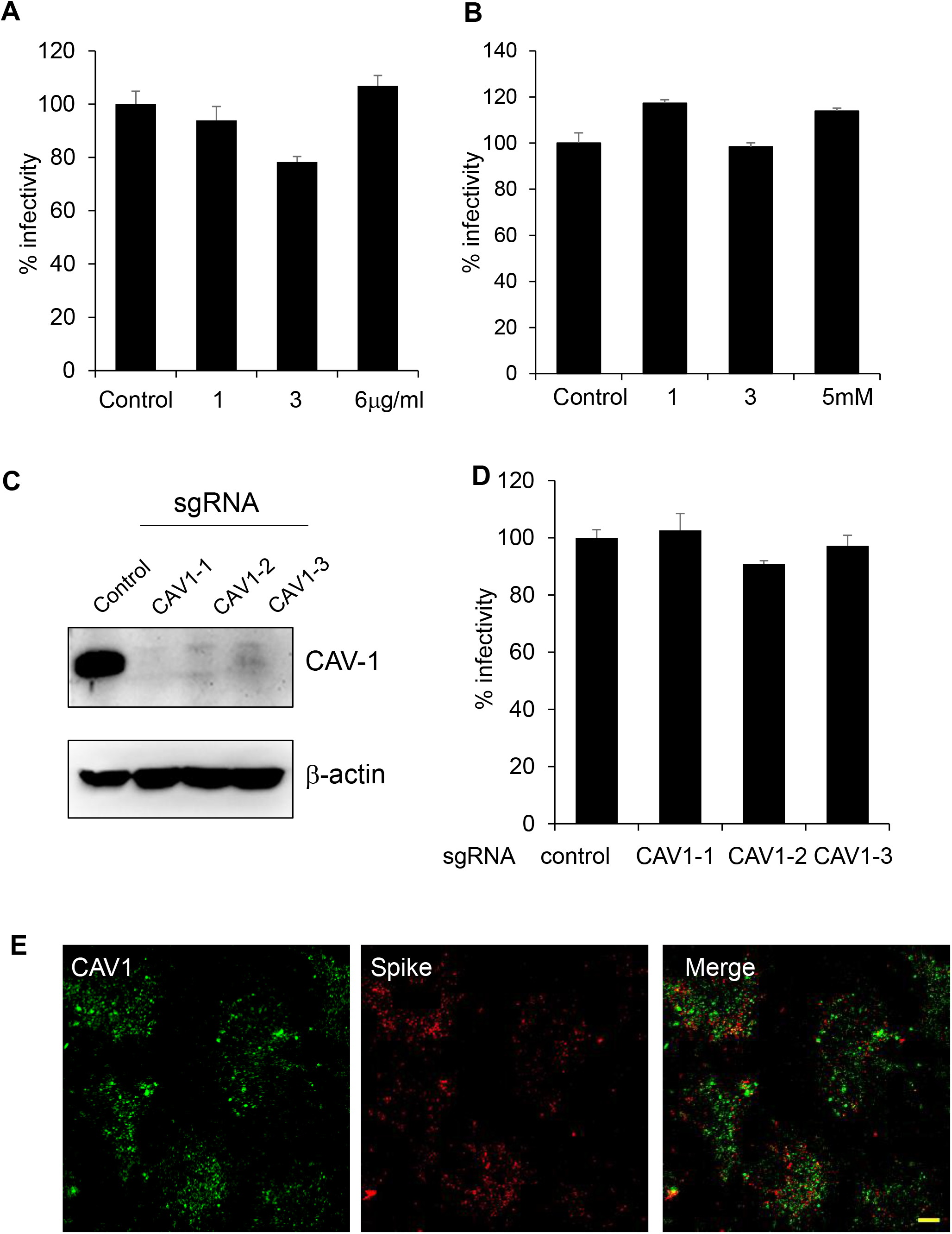
S pseudovirus transduces Huh 7 cells independent of caveolae-mediated endocytosis. A-B. Huh 7 cells pretreated with indicated concentrations of filipin or MβCD were transduced with S pseudovirus and luciferase were measured. Values are normalized to control and expressed as mean ± SD. C. Immunoblotting of CAV-1 expression in Huh 7 cells stably expressing a negative control or three independent CAV-1 sgRNAs with Cas9. D. Huh 7 cells as treated in (C) were transduced with S pseudovirus and luciferase were measured. Values are normalized to control and expressed as mean ± SD. E. Immunostaining of S pseudovirus infected Huh 7 cells for Spike(*red*) and CAV1(*green*). Bar, 10μm.

To further study whether the caveolae-dependent pathway was involved in the endocytosis of this virus, we knocked out the expression of caveolin-1(CAV-1, Figure 3C), which is the main scaffolding protein of the caveolae membranes in most cell types(31). Figure 3D showed that S pseudovirus could infect Huh 7 cells in the absence of CAV-1, suggesting the viral entry is independent of caveolae. This finding was further corroborated by the immunostaining experiment. The infected cells immune-labelled for SARS-CoV-2 spike protein and for caveolin-1 showed little colocalization (Figure 3E).

Taken together, these results indicated that SARS-CoV-2 is able to enter cells in a caveolae-independent manner.

### PI4P and PI(4,5)P_2_ are crucial for SARS-CoV-2 entry

Phosphoinositides (PI) are known to be involved throughout the process of endocytosis. Of note, PI(4,5)P_2_, PI4P and PI(3,4)P_2_ are essential molecules in endocytosis. We found that overexpression of either inositol polyphosphate-5-phosphatase E (INPP5E), which converts PI (4,5) P_2_ to PI4P, or the sac1 phosphatase, which dephosphorylates PI4P (Fig. 4A, Fig. S4), inhibited S pseudovirus transduction of 293-hACE2 cells (Fig. 4B), suggesting that PI4P and PI (4,5)P_2_ are involved in SARS-CoV-2 entry. Importantly, SARS-CoV entry has been reported to be dependent on phosphatidylinositol 4-kinase IIIβ(PI4KB)(23), and pretreating Huh 7 cells with phosphatidylinositol kinase inhibitors, PIK93 or wortmannin, inhibited S pseudovirus transduction at concentrations that effectively inhibited PI4-kinase activity (Fig. 4C), suggesting that PI4 kinase is also involved in SARS-CoV-2 infection. To better characterize the role of PI4KB in virus entry, we use shRNAs to knockdown PI4KB expression and found decreased virus infection in PI4KB knockdown cells (Fig.4D), indicating the role of PI4KB during viral entry.

### SARS-CoV-2 entry does not require the intracellular domain and catalytic sites of ACE2

Receptor-mediated endocytosis involves the interaction of the cytoplasmic tails of the receptor proteins with intracellular proteins to initiate internalization(32–34), and studies have shown that cytoplasmic domain of cell surface receptors could control endocytosis (35, 36). ACE2 is a type I transmembrane protein with a short cytoplasmic tail(7, 8). We therefore tested whether the cytoplasmic tail of ACE2 is required for SARS-CoV-2 entry. 293T cells overexpressing ACE2 mutant lacking the intracellular tail could be infected by SARS-CoV-2 pseudovirus as efficiently as wild-type ACE2 (Figure 5A-C). We further examined the receptor activity of ACE2 mutants with either the transmembrane domain of EGFR or the transmembrane domain and cytoplasmic tail of human ACE, the close homologue of ACE2 which could not support virus infection. Both mutants could support infection of SARS-CoV-2(Figure 5A-C). These results suggested that the cytoplasmic domain of ACE2 is not required for its SARS-CoV-2 infection and that there is no specificity of the transmembrane domain for its receptor activity.

**Fig 4.**
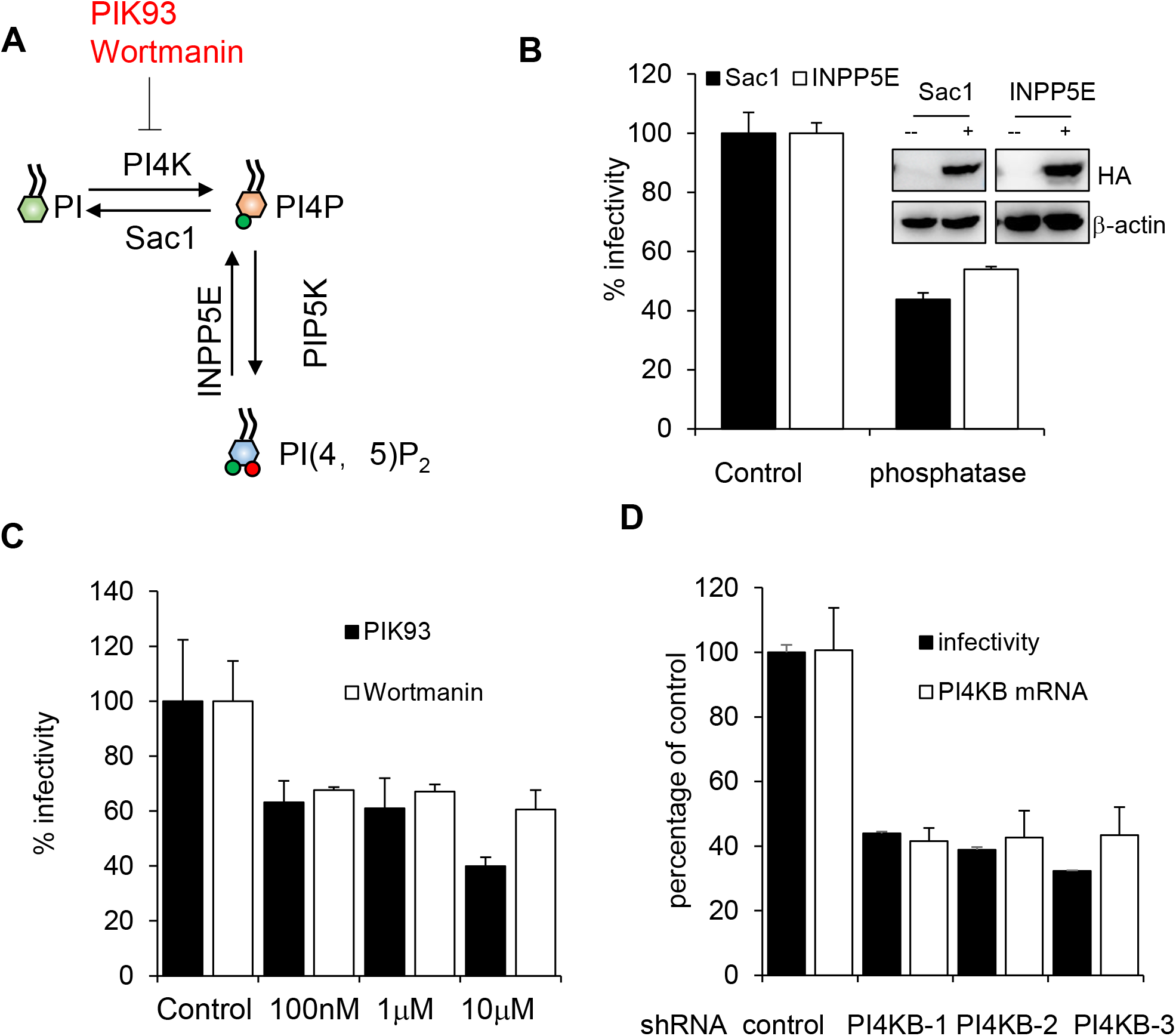
PI4KB is relevant to S pseudovirus infection. A. Diagram of phosphoinositide metabolic pathways. PIK93 and wortmannin were used as inhibitors targeting PI4K. B. 293T-ACE2 cells were transfected with HA-Sac1 or HA-INPP5E before they were transduced with S pseudovirus and luciferase were measured. Values are normalized to control and expressed as mean ± SD. Inset, immunoblotting of HA-Sac1 or HA-INPP5E expression with anti-HA antibody. C. Huh 7 cells pretreated with indicated concentrations of PIK93 or wortmannin were transduced with S pseudovirus and luciferase were measured. Values are normalized to control and expressed as mean ± SD. D. Huh 7 cells transduced with the indicated shRNAs were infected with S pseudovirus and luciferase were measured to reflect viral infection. PI4KB mRNA was quantified by qRT-PCR. Values are normalized to control shRNA.

**Fig 5.**
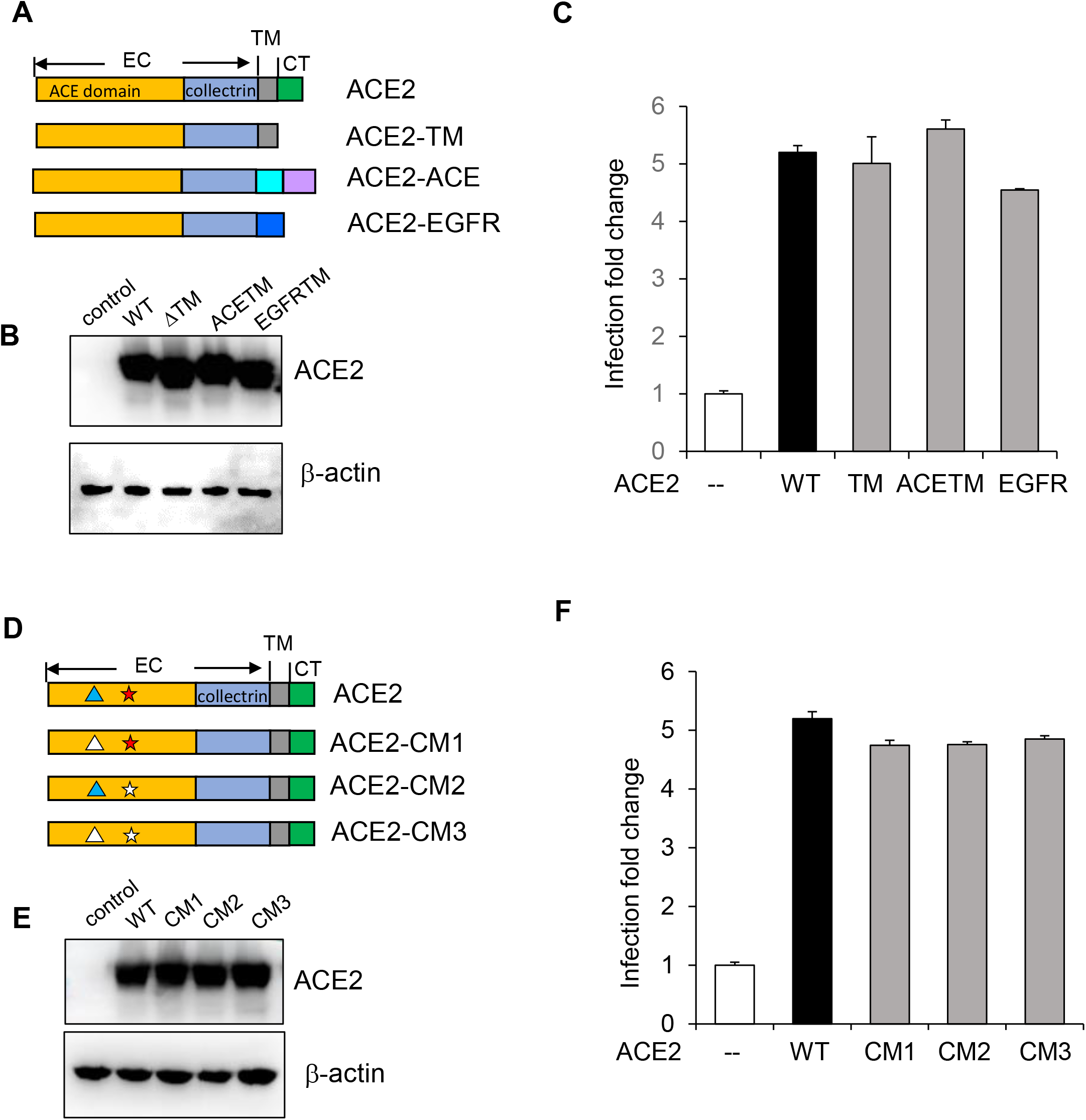
Cytoplasmic domain and enzymatic activity of ACE2 is not required for S pseudovirus infection. A. Diagram of ACE2 and ACE2 mutants. EC, extracellular domain. TM, transmembrane domain. CT, cytoplasmic tail. ACE2-TM, ACE2 lacking the cytoplasmic tail; ACE2-ACE, chimera of ACE2 with transmembrane and cytoplasmic tail form ACE; ACE2-EGFR, chimera of ACE2 with transmembrane domain form EGFR. B. Immunoblotting of ACE2 and ACE2 mutants expression. C. 293T cells transfected with wild type or indicated ACE2 mutant were transduced with S pseudovirus and luciferase were measured. Values are expressed as fold change to 293T cells. D. Diagram of ACE2 and ACE2 catalytic dead mutants. “∆” represents the Zinc binding histidines and “★” indicates the catalytic active site histidine. E. Immunoblotting of ACE2 and mutants expression. F. 293T cells transfected with wild type or indicated ACE2 mutant were transduced with S pseudovirus and luciferase were measured. Values are expressed as fold change to 293T cells.

We next tested whether the enzyme activity of ACE2 is required for SARS-CoV-2 infection. ACE2 is a Zinc metalloprotease with a single HEXXH zinc-binding motif(37, 38). In addition, ACE2 structure modeling and site-directed mutagenesis analysis showed that His505 plays an essential role in the catalysis of ACE2(39–41) (Fig 5D). To test whether enzymatic activity of ACE2 is required for SARS-CoV-2 entry, 293T cells overexpressing ACE2 that bears two zinc-binding histidine mutations (H374N&H378N, CM1) were transduced with S pseudovirus. Figure 5F showed that this mutant support viral infection equally well to wild type ACE2. Similarly, 293T cells overexpressing ACE2 catalytic site histidine mutation (H505L, CM2), or the combination of all three histidine mutations (H374N&H378N&H505L, CM3) could be transduced by S pseudovirus as efficiently (Fig.5D-F). These results suggested that the enzymatic activity of ACE2 is not required for SARS-CoV-2 infection.

## Discussion

Viral entry represents the first step of virus-host interaction and glycoprotein-bearing pseudoviruses have been wildly used for viral entry studies. Here we generated Spike protein bearing pseudoviruses that had either a luciferase or GFP reporter gene. These viruses could transduce host cells expressing ACE2 receptor and therefore could be used for viral entry studies.

For enveloped viruses, the viral envelop must fuse with host membranes either at cell surface or in endosomes before the successful release of viral genome into host cells. SARS-CoV and MERS-CoV have been reported to utilize both pathways depending on the availability of exogenous and membrane bound proteases(12). The entry of SARS-CoV-2 has been reported to require cell surface protease TMPRSS2(3), suggesting direct membrane at cell surface is possible. In addition, several studies have demonstrated that lysosomotropic agents could block SARS-CoV-2 entry(3, 4), indicating the virus may also enter via endocytosis. Here we got similar results when cells were treated with lysosomotropic agents. In addition, we found colocalization of spike protein with early endosome marker EEA1, further proved that this virus could enter cells via pH-dependent endocytosis.

As the best characterized endocytosis pathways, clathrin-mediated endocytosis has been reported to be involved in various viral infection, including SARS-CoV(14). Here by using different chemical inhibitors and CRISPR/Cas9 targeting the major protein in clathrin coat formation, we demonstrated that SARS-CoV-2 entry also was dependent on clathrin. While we are preparing this manuscript, Bayati et al. reported that the Spike protein of SARS-CoV-2 endocytosis is dependent on clathrin and S pseudovirus can be blocked by CHC siRNA treatment(42). Consistently, we also showed that SARS-CoV-2 pseudovirus enters cell via CME, in addition, we also showed that this process is independent of caveolae-mediated endocytosis, another common endocytosis pathway. The fact that Caco2 cells, which do not express caveolin (43), could support SARS-CoV-2 infection efficiently(44) also suggested that this virus could infect cells independent of caveolae.

PI4P and PI4KB have been reported to be essential for SARS-CoV infection(23) and a recent report showed that PIKfyve inhibitors could block S pseudovirus infection (4), which all suggested that phosphoinositides pathways are important for coronavirus infection. Here we further demonstrated that PI4P and PI(4,5)P_2_ are essential molecules for SARS-CoV-2 infection and PI4KB is also required for SARS-CoV-2 entry.

The role of cytoplasmic tail of ACE2 in SARS-CoV infection is controversial (14, 45). We found that the cytoplasmic domain as well as the enzymatic activity of ACE2 are not required for efficient virus entry, which are consistent with the facts that the receptor-binding domain of S interacts with the N-terminal peptidase domain of ACE2 and the enzymatic sites are not in direct contacts with spike as revealed by the S-ACE2 complex structure(46, 47).

In summary, we demonstrated that SARS-CoV-2 pseudovirus infects cells via pH-dependent endocytosis, which is dependent on clathrin, but not caveolae. Phosphoinositides pathways, particularly, PI4KB is required for efficient virus infection. In addition, we showed that the intracellular domain and the catalytic activity of ACE2 are not required for viral infection. Overall, these results provided new insights into SARS-CoV-2 cellular entry and present more precise targets for antiviral drug development.

## Materials and Methods

### Cell lines and constructs

Huh 7, Vero E6 and 293T cells were grown in Dulbecco’s modified Eagle medium (DMEM) supplemented with 10% fetal bovine serum (FBS), nonessential amino acids, 100 U/mL of penicillin, and 100 µg/mL of streptomycin.

ACE2, Sac1 and INPP5E were amplified from cDNA library prepared with Huh 7 cells and each was cloned into a lentiviral expression vector pSMPUW. ACE2 truncation and mutants were obtained with overlap extension PCR amplification using primers listed in Table S1. Codon-optimized spike of SARS-CoV-2 was purchased from GenScript and subcloned to pcDNA3.1. All constructs were confirmed by sequencing.

293T-ACE2 stable cell lines were generated by transfecting 293T cells with pSMPUM-ACE2 and selected with blasticidin. Antibiotic-resistant single clones were screened with immunoblotting and high ACE2-expressing clones were expanded for future use.

### Antibodies and reagents

Antibodies used in this study include: SARS-CoV-2 spike (GeneTex, Taiwan), HA tag, EEA1 (Cell Signaling Technology, Danvers, MA), HIV-p24 (SinoBiological, Beijing, China), β-actin (ABclonal, Woburn, MA), CHC (Santa Cruz Biotechnology, Dallas, TX), CAV1(Proteintech, Chicago, IL) and ACE2 (Abways Technology, Shanghai, China). PI4P and PI(4,5)P_2_ antibodies were purchased from Echelon Biosciences.

DAPI, Alexa Fluor conjugate CTB and Alexa Fluor conjugated secondary antibodies for microscopy experiments were purchased from Life Technologies.

Chloroquine, ammonium chloride, bafilomycin A1, chlorpromazine(CPZ), Filipin, MβCD were purchased from Sigma. Pitstop was purchased from Abcam. PIK93, Wortmannin were purchased from MedChemExpress.

### Pseudovirus preparation

Pseudovirus was prepared with a previously described protocol(4) with minor modification. Briefly, 293T cells were co-transfected with lentiviral packaging plasmid psPAX2, pcDNA3.1-Spike, pMD2.G, or empty vector, and lentiviral expression vector pSMPUW engineered with either GFP or NanoLuc luciferase. At 48h, 68h post-transfection, virus supernatants were harvested and filtered through 0.45 μm pore-size filters. Viral supernatants were then centrifuged at 30,000rpm for 2h in Himac ultracentrifuge S52ST rotor at 4 °C over a 20% sucrose solution, and virus stocks were resuspended with PBS and frozen at –80 °C.

### Pseudovirus entry and inhibition study

Unless otherwise indicated, Huh 7 cells were used for entry assays. For phosphatase inhibition test, 293T-ACE2 cells were first transfected with Sac1 or INPP5E. 30 hrs later, cells were split for immunoblotting and pseudovirus transduction. For ACE2 mutants assay, 293T cells were transfected with ACE2 or the indicated ACE2 mutant and 30 hrs later, cells were split for immunoblotting and pseudovirus transduction. To transduce cells with pseudovirus, cells seeded in 24-well plate were inoculated with 30ul of concentrated pseudovirus. About 36 hrs post inoculation, cells were lysed and NanoLuc luciferase activity was measured with Nano-Glo® Luciferase Assay System (Promega, Madison, WI) according to manufacturer’s instructions using a Synergy Neo 2 plate reader (BioTek, Winooski, VT). For cells transduced with GFP pseudovirus, photos were taken at 56 hrs post inoculation. For virus entry inhibition studies, cells were pretreated with indicated concentrations of inhibitors for 1 hr at 37 °C before pseudovirus inoculation. NanoLuc luciferase activity was measured as described above.

### Immunofluorescence staining

Immunofluorescence staining was performed as described in(48, 49). Images were taken with a Nikon C2 laser scanning confocal microscope in sequential scanning mode to limit crosstalk between fluorochromes.

### shRNA knockdown or CRISPR knockout cell lines construction

VSV-G pseudotyped shRNA or CRISPR lentiviruses were produced as previously reported (48, 50). Briefly, 293T cells were co-transfected with psPAX2, pMD2.G and pLKO.1-Puro based shRNA or pLentiCRISPR V2 based sgRNA lentiviral vectors and supernatants were harvested and used for transduction. sgRNA sequences and shRNA sequences used were listed in Table S2. Cells transduced with lentiviral particles were then selected with puromycin for stable cell lines.

### Statistics

All values represent means ± standard deviations and represent the results of a minimum of three independent experiments. Where applicable, the two-tailed Student’s t test was used to compare the means of control and experimental groups.

## Acknowledgements

This work was supported by the National Natural Science Foundation of China (81871662), Xi’an Jiaotong University Fund (xzy012019066 and xzy032020037) and Xi’an Jiaotong University Health Science Center-Qinnong Bank Fund (QNXJTU-04& QNXJTU-07).

